# Integrative analysis of Paneth cell proteomic and transcriptomic data from intestinal organoids reveals functional processes dependent on autophagy

**DOI:** 10.1101/410027

**Authors:** Emily J Jones, Zoe J Matthews, Lejla Gul, Padhmanand Sudhakar, Agatha Treveil, Devina Divekar, Jasmine Buck, Tomasz Wrzesinski, Matthew Jefferson, Stuart D Armstrong, Lindsay J Hall, Alastair JM Watson, Simon R Carding, Wilfried Haerty, Federica Di Palma, Ulrike Mayer, Penny P Powell, Isabelle Hautefort, Tom Wileman, Tamas Korcsmaros

**Affiliations:** Earlham Institute, Norwich Research Park, Norwich, NR4 7UZ, UK; Quadram Institute, Norwich Research Park, Norwich, NR4 7UA, UK; Norwich Medical School, University of East Anglia, Norwich, NR4 7TJ, UK; National Institute of Health Research, University of Liverpool, Liverpool, L3 5RF, UK; School of Biological Sciences, University of East Anglia, Norwich, NR4 7TJ, UK

**Keywords:** Paneth cells, Atg16l1, intestinal organoids, quantitative proteomics, selective autophagy

## Abstract

Using an integrative approach encompassing intestinal organoid culture, proteomics, transcriptomics and protein-protein interaction networks, we list Paneth cell functions dependent on autophagy.

**Abstract:** Paneth cells are key epithelial cells providing an antimicrobial barrier and maintaining integrity of the small intestinal stem cell niche. Paneth cell abnormalities are unfortunately detrimental to gut health and often associated with digestive pathologies such as Crohn’s disease or infections. Similar alterations are observed in individuals with impaired autophagy, a process which recycles cellular components. The direct effect of autophagy-impairment on Paneth cells has not been analysed. To investigate this, we generated a mouse model lacking *Atg16l1* specifically in intestinal epithelial cells making these cells impaired in autophagy. Using 3D intestinal organoids enriched for Paneth cells, we compared the proteomic profiles of wild-type (WT) and autophagy-impaired organoids. We used an integrated computational approach combining protein-protein interaction networks, autophagy targeted proteins and functional information to identify the mechanistic link between autophagy-impairment and disrupted pathways. Of the 284 altered proteins, 198 (70%) were more abundant in autophagy-impaired organoids, suggesting reduced protein degradation. Interestingly, these differentially abundant proteins comprised 116 proteins (41%), predicted targets of the selective autophagy proteins p62, LC3 and ATG16L1. Our integrative analysis revealed autophagy-mediated mechanisms degrading proteins key to Paneth cell functions, such as exocytosis, apoptosis and DNA damage repair. Transcriptomic profiling of additional organoids confirmed that 90% of the observed changes upon autophagy alteration affect protein level and not gene expression. We performed further validation experiments showing differential lysozyme secretion, confirming our computationally inferred down-regulation of exocytosis. Our observations could explain how protein level alterations affect Paneth cell homeostatic functions upon autophagy impairment.

## Introduction

Paneth cells, located at the bottom of the crypts of Lieberkühn in the small intestine, secrete various types of antimicrobial compounds (e.g. lysozyme, defensins) regulate the microbial composition of the intestine as well as growth factors that maintain the crypt-associated stem cell population(Bevins and Salzman, 2011). The secretory activity of Paneth cells strongly relies on pathways to release proteins like antimicrobials into the lumen as one of the gut barrier functions contributing to intestinal homeostasis (Bel et al., 2017; Liu et al., 2013). Conventional protein secretion involves trafficking through ER and Golgi(Farquhar and Palade, 1981; Viotti, 2016). Paneth cell defects such as altered granule morphology and increased susceptibility to ER stress are seen in mouse models where autophagy is lost from intestinal epithelial cells (Liu et al., 2013; Wileman, 2013).

Autophagy is a pivotal recycling process that sequesters cytoplasmic misfolded proteins or damaged organelles as well as clears the cytosol from invading pathogens. These targets are captured in double-membrane vesicles called autophagosomes that are subsequently delivered for degradation to lysosomal compartments (Deretic et al., 2013; Glick et al., 2010; Todde et al., 2009; Wileman, 2013). Although initially considered as a non-selective process elicited upon starvation, stress or infection, recent studies have indicated that the cargoes of autophagy, be it organelles (such as mitochondria, peroxisomes, ribosomes, endoplasmic reticulum), pathogens or protein aggregates, are recognized in a very selective manner, termed selective autophagy (Fimia et al., 2013; Zaffagnini and Martens, 2016). Sequestration of selective autophagy-targets follows recognition by specific cargo receptors and involves the Atg12-Atg5-Atg16 complex, instrumental in the early stages of the autophagosome biogenesis by determining the site of LC3 lipidation (Fujita et al., 2008). Through LIR (LC3 interacting region) motifs, the lipidated LC3 adaptor not only targets various cargoes for sequestration but also recruits multiple autophagy receptor proteins such as p62, NDP52, NBR1, NIX and Optineurin. Cargo recognition by the autophagy receptors happens generally *via* ubiquitin-dependent or ubiquitin-independent mechanisms (Khaminets et al., 2016). The autophagy receptors bridge their cargoes (which are specifically targeted by the presence of receptor recognition motifs and degradation signals) with the autophagosomal membrane (Stolz et al., 2014). These events eventually result in the cargo engulfment by the autophagosome, which then fuses with the lysosome to form the autophagolysosome in which the contents are degraded by lysosomal enzymes (Glick et al., 2010; Todde et al., 2009). In addition to its recycling role, autophagy is involved in Paneth cell response to microbial exposure. Upon microbial challenge, lysozyme secretion by Paneth cells is conducted through the diversion of degradative autophagy towards a secretory process, named secretory autophagy pathway (Bel et al., 2017; Kimura et al., 2017), although various autophagy-independent secretory pathways have also been reported (Barlowe and Miller, 2013).

Alteration in secretory autophagy have been associated with many intestinal diseases. Severe gut pathologies such as Crohn’s disease (CD), an inflammatory bowel disease (IBD) or food-borne pathogen infections (e.g. Salmonellosis) are associated with Paneth cell dysfunctions including disrupted antimicrobials production as observed in chronic inflammatory and infectious diseases (Liu et al., 2016; Martinez Rodriguez et al., 2012; Perminow et al., 2010; Salzman et al., 2003; Wehkamp et al., 2005). GWAS studies have identified mutations in autophagy-related genes, in particular, mutation in the key autophagy gene *ATG16L1* resulting in granule exocytosis abnormalities in Paneth cells with a negative effect on autophagy-mediated defense against bacterial pathogens (Cadwell et al., 2008; Lassen et al., 2014; Perminow et al., 2010; Wehkamp et al., 2005). Due to its critical function in the autophagy machinery, ATG16L1 is required for the proper functioning of autophagy in general (Kuballa et al., 2008; Mizushima et al., 2003) and in various intestinal cell types, including Paneth cells (Cadwell et al., 2008; Patel et al., 2013). In mice harbouring mutations in key autophagy genes such as *Atg7* or *Atg16l1*, lysozyme levels were decreased, granule size was reduced, and exocytosis was abnormal in Paneth cells compared with wild-type mice (Cadwell et al., 2008; Conway et al., 2013; Lassen et al., 2014; Wittkopf et al., 2012). Specific mutations in *Atg16l1* such as T300A affect the activity of *Atg16l1* due to the gain of a Caspase 3 cleavage site without compromising the protein architecture(Salem et al., 2015). Even though the critical role of ATG16L1 in modulating autophagy in Paneth cells is known, the exact molecular mechanisms and cellular processes affected by autophagy-impairment remain to be elucidated.

In this study, we use the small intestinal organoid culture model, which reproduces crypt-like and villus-like domains characteristic of intestinal morphology recapitulating many functions of the small bowel. Intestinal organoids contain specialised cell types such as Paneth cells that cannot be examined in cell lines making them a unique model system to analyse Paneth cell proteins and functions (Sato et al., 2009). To increase the usefulness of the organoid model, we enrich both WT and autophagy-impaired organoids for Paneth cells by directing the lineage of organoid differentiation by following established protocols (see Methods section). Thereafter, we analyse the quantitative proteome of Paneth cell-enriched small intestinal organoids specifically lacking *Atg16l1* in intestinal epithelial cells, and compare it to the proteomic profile of WT Paneth cell-enriched organoids. Given the known defects of autophagy in inflammatory disorders, the major autophagy impairment due to the loss of Atg16l1 could be considered as an extreme disease model. In order to understand the possible mechanisms by which autophagy impairment could modulate the abundance of proteins in key epithelial cell functions, we establish an *in-silico* workflow (**Figure 1**) combining several computational approaches including protein-protein interaction networks, interaction evidence incorporating protein targeting by selective autophagy and information on functional processes. Using this integrative approach, we show that proteins with altered abundances in the autophagy-impaired Paneth cell-enriched organoids, could be substrates of selective autophagy and could be targeted by autophagy resulting in their degradation. Our integrative approach pointed out several autophagy-dependent cellular processes as well as novel mechanisms in which autophagy was influencing those processes. With the transcriptomics profiling of the WT and autophagy impaired organoids we validate that the proteomic changes are due to protein level alterations and not due to gene expression changes. Importantly, we also confirm that autophagy dysfunction alters several cellular processes such as cellular exocytosis which was downregulated in autophagy-impaired organoids and is known to be deleteriously altered in patients with inflamed digestive tract (e.g. CD patients). Taken together, our observations based on a model of autophagy impairment in Paneth cells provides a mechanistic explanation of Paneth cell dysfunction due to autophagy-impairment. The demonstrated involvement of novel autophagy-dependent processes in Paneth cells extends our understanding of disorders related to autophagy dysfunction. Furthermore, it opens the door for the development of new and/or supplementary therapeutic interventions for digestive pathologies triggered or exacerbated upon autophagy deficiency.

**Figure 1.**
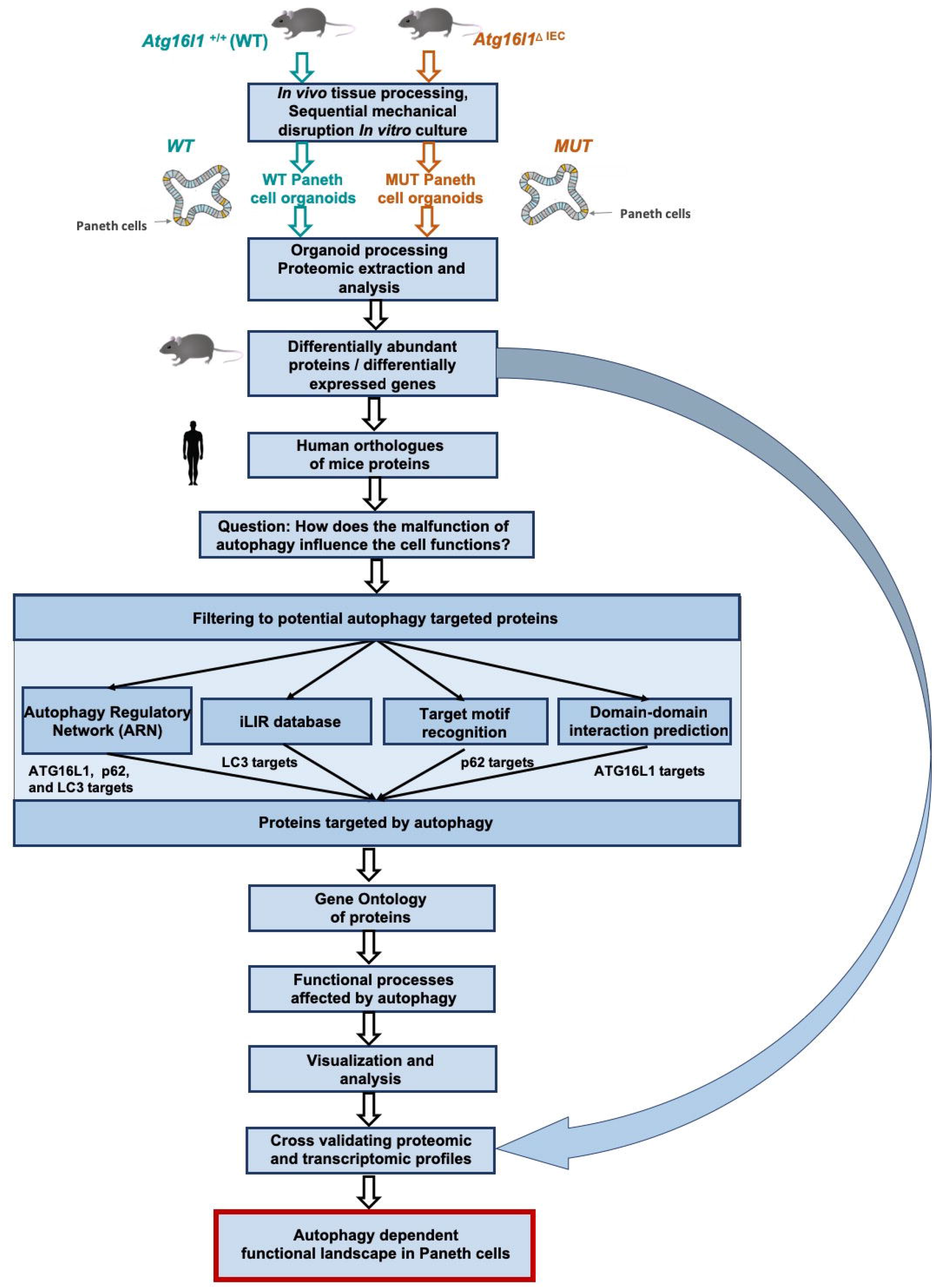
A schematic representation of the workflow to determine the functional effects of the autophagy impairment in *Atg16l11*^⟀IEC^ in Paneth cell organoids. Three biological replicates were generated for each condition and genotype tested and for both proteomics and transcriptomics type of profiling.

## Methods

### Animal handling

C57/Bl6 mice of both sexes were used for organoid generation. All animals were maintained in accordance with the Animals (Scientific Procedures) Act 1986 (ASPA).

### Generation of *Atg16l1*^flox/−VilCre^ (*Atg16L1*^⟀IEC^) mice

Mice were generated using a Cre/LoxP system. Briefly, LoxP sites were inserted either side of *Atg16l1* exon 2, creating *Atg16l1*^flox/+^ mice. Crossing these mice with PGK-Cre mice, expressing Cre recombinase under the PGK promoter that is in all cell types, led to the excision of the floxed exon 2 in one allele by Cre recombinase. This in turn introduced a stop codon, generating *Atg16l1*^+/−^ mice. Cell type-specific *Atg16l1* deletion was induced using a Villin promoter to drive expression of Cre recombinase only in villin-expressing cells. This was achieved by crossing *Atg16l1*^flox/flox^ mice with *Atg16l1*^+/−VilCre^ mice to produce *Atg16l1*^flox/−VilCre^ mice deficient in *Atg16l1* in intestinal epithelial cells. *Atg16l1*^Fl/+^ mice were used as wild-type controls (WT). Transgenic mice were genotyped using end-point PCR and gel electrophoresis. *Atg16l1* alleles were firstly designated as WT (+), floxed (FI) or knock-out (−) and the presence of the *Cre* recombinase gene under the control of the *villin* promoter subsequently designated positive (Vil-Cre) or negative. Combining the PCR results identified the *Atg16l1^Fl/+^* (WT) or *Atg16l1^FI/−VilCre^* (*Atgl6L1*^⟀*IEC*^ KO) mice. All primers used to validate the organoid models are listed in **Supplementary table 1**.

### Small intestinal organoid cultures for both proteomics and transcriptomics profiling

Murine small intestinal organoids were generated as described previously (Sato et al., 2013). Briefly, the entire small intestine was opened longitudinally, washed in cold PBS then cut into ~5mm pieces. The intestinal fragments were incubated in 30mM EDTA/PBS for 5 minutes, transferred to PBS for shaking, then returned to EDTA for 5 minutes. This process was repeated until five fractions were generated. The PBS supernatant fractions were inspected for released crypts. The crypt suspensions were passed through a 70μm filter to remove any larger villus-containing fragments, then centrifuged at 300xg for 5 minutes. For 3D organoid cultures, pellets were resuspended in 200μl phenol red-free Matrigel (Corning), seeded in small domes into 24 well-plates and incubated at 37⟀C for 20 minutes to allow Matrigel to polymerise. Organoid medium (Advanced DMEM/F12 (Life Technologies)) containing growth factors including EGF (50 ng/ml, Life Technologies), Noggin (100 ng/ml, PeproTech) and R-spondin 1 (500 ng/ml, R&D) was then overlaid. For 2D organoid monolayers used for the lysozyme secretion validation, pellets were resuspended in organoid media and overlaid onto coverslips coated with phenol red-free Matrigel (Corning). For the quantitative proteomics analysis, the Paneth cell population in 3D WT and *Atg16l1^ΔIEC^* organoids were enriched by addition of 3μM CHIR99021 (Tocris) and 10μM DAPT (Tocris) to the organoid culture media on day 2, 5 and 7 post crypt isolation according to previously published and well established protocols(Nakanishi et al., 2016; Yin et al., 2014). For both proteomics and transcriptomics profiling, replicate organoids were generated from three individual animals for each genotype and each enrichment condition tested, generating three biological replicates for each sample group. Two additional biological replicates of both WT normally differentiated, and Paneth cell-enriched organoids were also generated, profiled by proteomics and used as controls to validate the Paneth cell enrichment. One replicate was subsequently removed from the WT Paneth cell-enriched transcriptomics dataset as initial analysis showed it as an outlier.

### qPCR, Immunoblotting and Western blots

To confirm WT and *Atg16l1*^ΔIEC^ organoids retained the intestinal phenotype and expressed the intestinal epithelial cell-type markers, gene expression was analysed by qPCR. On day 8 post crypt isolation, normally differentiated organoid pellets were lysed in 500μl TRIzol (Life Technologies). RNA was extracted by chloroform extraction followed by precipitation in isopropanol and ethanol. Superscript II reverse transcription protocol was used with 200ng organoid RNA to generate single stranded cDNA. Gene specific primers for *Lgr5* (stem cells), *Villin* (epithelial cells), *Chromogranin A* (enteroendocrine cells), *Mucin 2* (goblet cells), *CD24* (Paneth cells) and *β-actin* were used for linear amplification of cDNAs using limited number of cycles. B-actin was used as a housekeeping gene expression analysis. Primers and PCR cycle numbers are listed in **Supplementary table 1**. To confirm the absence of the Atg16l1 protein, impairment in autophagy and lysozyme quantification, specific protein expression was analysed by immunoblotting and Western blot analysis. On day 8 post crypt isolation, normally differentiated or Paneth enriched organoid pellets were lysed in m-PER lysis buffer (ThermoFisher Scientific) containing protease inhibitor cocktail (Roche). 3-30μg protein per well was separated using NuPAGE precast 4-12% polyacrylamide gel system (ThermoFisher Scientific). Immunoblotting was carried out using an X-Cell II blot module (ThermoFisher Scientific) onto polyvinylidene (PVDF) membranes (ThermoFisher Scientific). Membranes were probed for ATG16L1 (Abgent #AP1817b), LC3II (Sigma #L7543), β-actin (Sigma #A1978 (Clone AC015)) and lysozyme (Dako #A0099 EC3.2.1.17) and protein bands visualized using Odyssey infrared imaging system (Li-Cor) at 700nm and 800 nm. Densitometry of lysozyme bands was performed using the FIJI/ImageJ package and expressed as arbitrary units (AU) from at least 3 biological replicates for each group.

### Protein sample preparation for proteomics

On day 8 post crypt-isolation, Paneth enriched 3D organoids were extracted from Matrigel using Cell Recovery Solution (BD Bioscience), washed in PBS and centrifuged at 300xg for 5 minutes. Organoid pellets were lysed by sonication in 1% (w/v) sodium deoxycholate (SDC) in 50mM ammonium bicarbonate. Samples were heated at 80°C for 15 min before centrifugation at 12,000xg to pellet debris. The supernatant was retained, and proteins reduced with 3 mM DTT (Sigma) at 60°C for 10 min, cooled, then alkylated with 9 mM iodoacetamide (Sigma) at RT for 30 min in the dark; all steps were performed with intermittent vortex-mixing. Proteomic-grade trypsin (Sigma) was added at a protein:trypsin ratio of 50:1 and incubated at 37°C overnight. SDC was removed by adding TFA to a final concentration of 0.5% (v/v). Peptide samples were centrifuged at 12,000 x g for 30 min to remove precipitated SDC.

### NanoLC MS ESI MS/MS analysis

Peptides were analysed by on-line nanoflow LC using the Ultimate 3000 nano system (Dionex/Thermo Fisher Scientific) system coupled to a Q-Exactive HF mass spectrometer (Thermo Fisher Scientific) essentially as described in (Dong et al., 2017). Peptides were separated by an Easy-Spray PepMap^®^ RSLC analytical column (50 cm × 75 μm inner diameter, C18, 2 μm, 100 Å) fused to a silica nano-electrospray emitter (Dionex). Column temperature was kept at a constant 35°C. Chromatography buffers consisted of 0.1 % formic acid (buffer A) and 80 % acetonitrile in 0.1 % formic acid (buffer B). The peptides were separated by a linear gradient of 3.8 – 50 % buffer B over 90 minutes at a flow rate of 300 nl/min. The Q-Exactive HF was operated in data-dependent mode with survey scans acquired at a resolution of 60,000. Up to the top 10 most abundant isotope patterns with charge states +2 to +5 from the survey scan were selected with an isolation window of 2.0Th and fragmented by higher energy collisional dissociation with normalized collision energies of 30. The maximum ion injection times for the survey scan and the MS/MS scans were 100 and 45ms, respectively, and the ion target value was set to 3E6 for survey scans and 1E5 for the MS/MS scans. MS/MS events were acquired at a resolution of 30,000. Repetitive sequencing of peptides was minimized through dynamic exclusion of the sequenced peptides for 20s.

### Protein identification and quantification

Spectral data was imported into Progenesis QI (version 4.1, Nonlinear Dynamics). Runs were time aligned using default settings and using an auto selected run as reference. Peaks were picked by the software using default settings and filtered to include only peaks with a charge state between +2 and +7. Spectral data were converted into .mgf files with Progenesis QI and exported for peptide identification using the Mascot (version 2.3.02, Matrix Science) search engine. Tandem MS data were searched against a database including translated ORFs from the *Mus musculus* genome (Uniprot reference proteome (reviewed), UP000000589, February 2017) and a contaminant database (cRAP, GPMDB, 2012) (combined 17,010 sequences; 9,549,678 residues. Precursor mass tolerance was set to 10 ppm and fragment mass tolerance was set as 0.05 Da. Two missed tryptic cleavages were permitted. Carbamidomethylation (cysteine) was set as a fixed modification and oxidation (methionine) set as a variable modification. Mascot search results were further validated using the machine learning algorithm Percolator embedded within Mascot. The Mascot decoy database function was utilised and the false discovery rate (FDR) was <1%, while individual percolator ion scores >13 indicated identity or extensive homology (p <0.05). Mascot search results were imported into Progenesis QI as XML files. Peptide intensities were normalised against the reference run by Progenesis QI and these intensities were used to highlight relative differences in protein expression between sample groups. Only proteins with 2 or more identified peptides were included in the dataset. Statistical analysis (one factor ANOVA) of the data was performed using Progenesis QI to identify significantly (p < 0.05, absolute relative fold change ≥ 2, number of unique peptides ≥ 2) proteins with altered abundances. The proteomic dataset has been submitted to PRIDE (accession id: PXD010940).

### RNA sample preparation for transcriptomics

For transcriptomic profiling, Paneth cell-enriched organoids were extracted from Matrigel on day 8, recovered in Cell Recovery Solution (BD Bioscience) and washed in PBS. RNA extraction was performed using Exiqon tissue kit according to the manufacturer’s protocol. Stranded RNA seq libraries were constructed using the NEXTflex™ Rapid Directional RNA-Seq Kit (PN: 5138-07) using the poly-A pull down beads from Illumina TruSeq RNA v2 library construction kit (PN: RS-122-2001) with the NEXTflex™ DNA Barcodes – 48 (PN: 514104) diluted to 6uM. RNA QC was carried out using Qubit DNA kit (Life technologies Q32854 & Q32852) and PerkinElmer GX RNA assay (PN:CLS960010) were prior to library construction. In more details, mRNAs were extracted with a poly-A pull down using biotin beads, fragmented and first strand cDNA was synthesised. This process reverse transcribes the cleaved RNA fragments primed with random hexamers into first strand cDNA using reverse transcriptase and random primers. The second strand synthesis process removes the RNA template and synthesizes a replacement strand to generate ds cDNA. Directionality is retained by adding dUTP during the second strand synthesis step and subsequent cleavage of the uridine containing strand using Uracil DNA Glycosylase. The ends of the samples were repaired using the 3’ to 5’ exonuclease activity to remove the 3’ overhangs and the polymerase activity to fill in the 5’ overhangs creating blunt ends. A single ‘A’ nucleotide was added to the 3’ ends of the blunt fragments to prevent them from ligating to one another during the adapter ligation reaction. A corresponding single ‘T’ nucleotide on the 3’ end of the adapter provided a complementary overhang for ligating the adapter to the fragment. This strategy ensured a low rate of chimera formation. The ligation of a number indexing adapters to the ends of the DNA fragments prepared them for hybridisation onto a flow cell. The ligated products were subjected to a bead-based size selection using Beckman Coulter XP beads (PN: A63880). As well as performing a size selection this process removed the majority of unligated adapters. Prior to hybridisation to the flow cell the samples were PCR’d to enrich for DNA fragments with adapter molecules on both ends and to amplify the amount of DNA in the library. The strand that was sequenced is the cDNA strand. The insert size of the libraries was verified by running an aliquot of the DNA library on a PerkinElmer GX using the High Sensitivity DNA chip (PerkinElmer CLS760672) and the concentration was determined by using a High Sensitivity Qubit assay and qPCR. Libraries were then equimolar pooled and checked by qPCR to ensure the libraries had the necessary sequencing adapters ligated.

### Sequencing the stranded RNA libraries

The constructed stranded RNA libraries were normalised and equimolar pooled, the final pool was quantified using a KAPA Library Quant Kit and found to be 2nM for the WT samples and 9.8nM for the Atg16l1ΔIEC samples. 9.5μl (WT) and 2.04μl (*Atg16l1*^ΔIEC^) of the pool was combined with 0.5μl 2N NaOH to make a 2nM dilution. This was incubated for five minutes at room temperature to denature the libraries before 990μl of HT1 was added to make a 20pM dilution. 60μl of the 20pM dilution was combined with 60μl of HT1 plus a 1% PhiX spike in (Illumina FC-110-3001) for each lane the pool was run in to make the final running concentration of 10pM. The flow-cell was clustered using HiSeq PE Cluster Kit v3 (Illumina PE-401-3001) for the WT samples and HiSeq PE Cluster Kit v4 (Illumina GD-401-4001) for the Atg16l1ΔIEC samples. The lllumina PE_HiSeq_Cluster_Kit_V3_cBot_recipe_V8.0 (WT) and PE_HiSeq_Cluster_Kit_V4_cBot_recipe_V9.0 (Atg16l1ΔIEC) methods were used on the Illumina cBot. Following the clustering procedure, the flow-cell was loaded onto the Illumina HiSeq2000 (WT) or HiSeq2500 (Atg16l1ΔIEC) instrument following the manufacturer’s instructions with a 101 cycle paired reads and a 7 cycle index read (WT) or a 126 cycle paired end reads and a 12bp/6bp dual index read (Atg16l1ΔIEC). For WT samples, the sequencing chemistry used was HiSeq SBS Kit v3 (Illumina FC-401-3001) with HiSeq Control Software 2.2.68 and RTA 1.18.66.3. For Atg16l1ΔIEC samples the sequencing chemistry used was HiSeq SBS Kit v4 (Illumina FC-401-4003) with HiSeq Control Software 2.2.58 and RTA 1.18.64. Reads in bcl format were demultiplexed based on the 6bp Illumina index by CASAVA 1.8, allowing for a one base-pair mismatch per library, and converted to FASTQ. format by bcl2fastq.

### Mapping and identification of differentially expressed transcripts

The quality of stranded reads was assessed by FastQC software (version 0.11.4). Gene and transcript abundances were estimated with kallisto (version 0.44.0) (Bray et al., 2016). The Sleuth R library was used to perform differential gene expression (0.30.0) (Pimentel et al., 2017). mRNAs and lncRNAs with an absolute log2 fold change of 1 and q value <= 0.05 were considered to be differentially expressed.

### Lysozyme activity assay

Lysozyme activity associated with Paneth cell antimicrobial defense was measured in 2D organoid culture medium using the EnzChekTM Lysozyme Assay Kit, according to manufacturer’s instructions (ThermoFisher Scientific). Briefly, 2D organoids were cultured from WT and *Atg16l1*^ΔIEC^ mice as described in the relevant method section. Following a 20 hours post-seeding incubation, the organoid culture medium was collected. Remaining cellular debris were removed by centrifugation at 600xg for 5 minutes and filtration on 0.20μm PES filters. FITC-fluorescence, proportional to the lysozyme activity released by Paneth cells into the medium, was measured on a Fluostar Optima Fluorometer (BMG Labtech), and corrected for background fluorescence. Lysozyme activity expressed as U/ml was determined from standard curves on at least 3 biological replicates for each group.

### Cross-validation of transcriptomic and proteomic profiles

To compare the quantity of gene expression and protein abundance, Ensembl gene IDs were converted to Uniprot IDs to unify the identities using the ID mapping tool of Uniprot. We defined a cut off value, if the difference between the log_2_ based fold change values (regarding the protein abundance and gene expression) is higher than 0.7, hypothetically the change in protein abundance level could happen due to the malfunction of autophagy.

### Interaction resources and computational methods to identify proteins targeted by autophagy

In order to make the proteomic data comparable to human interaction networks, the human orthologs of the proteins with altered abundances from the Paneth cell organoids were identified using InParanoid (Sonnhammer and Östlund, 2015). To identify the autophagy targeted protein components, the interaction partners of the three autophagy receptor and adaptor proteins, namely p62, LC3 and ATG16L1, were retrieved from the manually curated section of the Autophagy Regulatory Network (Türei et al., 2015). To enhance the coverage and improve interpretations, the interactions retrieved from experimental data were complemented with the predicted targets of p62, LC3 and ATG16L1. The putative targets of p62 and ATG16L1 were inferred using in-house custom scripts written in the Python programming language. The predictions of p62 and ATG16L1 targets were based on the standard motif search and domain-domain interaction prediction methods (Korcsmaros et al., 2013), respectively. The p62 recognition motif was retrieved from (Jadhav et al., 2011). For the domain-domain interaction prediction method, the known set of interacting domain-pairs were obtained from the DOMINE database (Raghavachari et al., 2008). DOMINE captures information about interacting domain-pairs from experiments, structural studies and predictions. Domain annotations for all proteins were retrieved from UniProt (The UniProt Consortium, 2017). The targets of LC3 were downloaded from the iLIR database (Jacomin et al., 2016).

### Functional analysis to identify affected processes

To check whether the changes are due to Paneth cell enrichment or because of the impaired autophagy we carried out a control experiment where we compared the proteomic profile of normal intestinal organoid to Paneth cell-enriched organoids. To assess the functional importance of proteins with altered abundances targeted by autophagy within a network context, Gene Ontology Biological Process terms (Ashburner et al., 2000) derived from UniProt (The UniProt Consortium, 2017) were used. Biological Process terms not related to the intestine (action potential, development of other organs/tissues etc) were discarded following a manual curation of the terms in order to maintain the intestinal context (**Supplementary table 2**). We performed an additional, extensive manual curation to determine the role of all the proteins in these biological processes as well as to define their effect (activational or inhibitory) on these processes. Based on the directionality of the interaction between p62/LC3/ATG16L1 and the proteins with altered abundances, we inferred the direction of modulation of the assigned biological process terms. The aggregated trends of the processes were inferred as follows: the processes were considered to be either up-regulated or down-regulated if more than 70% of its interactions were classified as stimulatory or inhibitory, respectively. In cases where the singular proportion is less than 70%, the functional process was considered to have dual modulation (i.e., both up- and down-regulated). The networks were visualized in Cytoscape 3.5.1 (Franz et al., 2016).

## Results

### Paneth cell-enriche *Atg16l1*1^⟀IEC^ organoids are a valid model to study the role of autophagy in intestinal epithelial homeostasis

We have generated a mouse model that lacks *Atg16l1* specifically in intestinal epithelial cells (*Atg16l1*^ΔIEC^) and have used self-organizing *in vitro* organoid cultures generated from small intestinal crypts (Sato et al., 2009) to assess the impact of autophagy on Paneth cell functions. As expected, normally differentiated organoids from both WT and *Atg16l1*^ΔIEC^ mice included viable budding crypts that expanded from a core organoid (**Figure 2A**). Detection of mRNA transcripts by linear qPCR for *Lgr5, ChgA, Muc2* and *Cd24* cDNAs along the housekeeping *β*-Actin gene revealed that *Atgl6l1*^ΔIEC^ organoids expressed markers for stem cells, enteroendocrine cells, goblet cells and Paneth cells at similar levels as WT organoids (**Figure 2B**), confirming similar differentiation expression levels in both genotypes of important cell types found in the *in vivo* small intestinal epithelium. We observed that the villin transcript shows a slight reduction in the KO organoids compared with the WT ones but remains indicative of the presence of enterocytes in both organoid models. In particular, we noted that the level of the Paneth cell marker CD24 was similar between WT and *Atg16l1*^ΔIEC^ organoids (**Figure 2B**), suggesting that the number of Paneth cells was similar in both genetic backgrounds.

**Figure 2.**
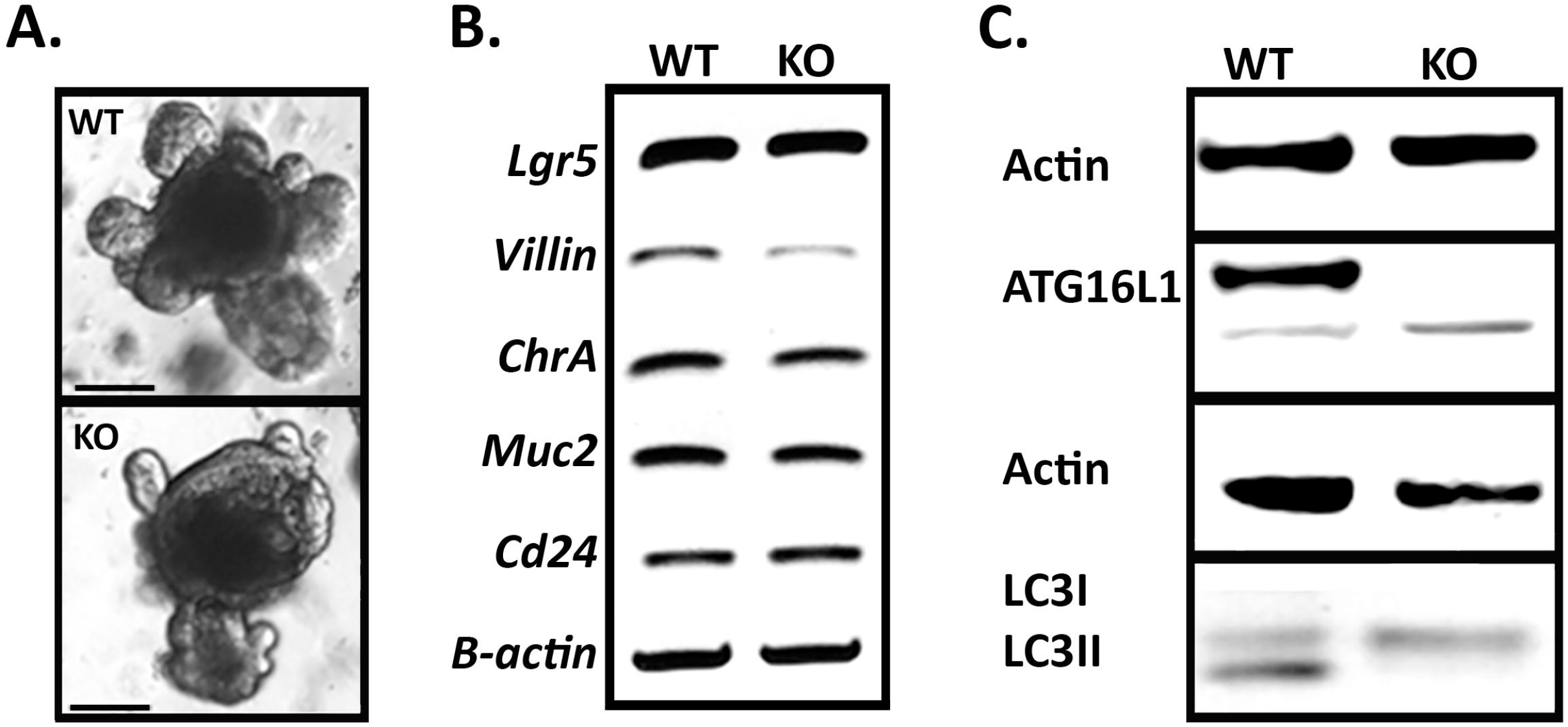
*Atg16l11*^⟀IEC^ organoids contain the same intestinal epithelial cell types as WT organoids but lack Atg16l1 and LC3II both in the transcriptional and protein levels (n=3). A) Brightfield image of control (WT) and *Atg16l1*^⟀IEC^ (KO) organoids after 7-days of growth. Magnification x10, scale bar 100μm. B) Cell type-specific qPCR amplification of cDNAs in WT and *Atg16l1*^⟀IEC^ (KO) organoids. *β-Actin* was used as a housekeeping gene. C) Western blots using anti-ATG16Ll and LC3 antibodies detected Atg16l1 and LC3II in control organoids but *Atg16l1*^⟀IEC^(KO) organoids were deficient in Atg16l1 and LC3II.

To increase the technical feasibility of investigating Paneth cells’ dependency on autophagy, organoids were further enriched for Paneth cells using a well-established and published protocol, presented in details in the Methods (Nakanishi et al., 2016; Yin et al., 2014). We confirmed Paneth cell enrichment using two complementary approaches. First, we observed the transcriptomics profiles generated from WT and *Atg16l1*^ΔIEC^ genotypes. For each genotype we generated a list of differentially expressed genes by comparing Paneth cell-enriched to normally differentiated organoids. Genes previously identified as Paneth cell markers were observed in these lists of differentially expressed genes (Haber et al., 2017). Paneth cell markers were significantly enriched in the gene expression dataset generated from WT organoids (56/83 markers present, hypergeometric test, p= 5.1e-57) and from *Atg16l1*^ΔIEC^ organoids (58/83 markers present, hypergeometric test, p= 3.2e-14), suggesting Paneth cells were well successfully enriched through the applied enrichment protocol. Second, we compared the proteomic profiles of normally differentiated intestinal organoids with that of Paneth cell-enriched organoids, focussing on organoids of WT and *Atg16l1*^ΔIEC^ genotypes (**Supplementary table 3, Supplementary table 4**). Using a similar workflow as for the subsequent comparison between the WT and Atg16l1-deficient Paneth cell-enriched organoids (**Figure 1**), we observed proteins with significantly different abundance between normally differentiated and Paneth cell-enriched organoids. We observed that Paneth cell specific processes were altered upon impaired autophagy in the enriched organoids. Proteins related to exocytosis, proteasome-ubiquitin system-related processes, immune response and apoptosis were differentially abundant in Paneth cell-enriched organoids. Overall, the two approaches support the conclusion that the organoids used in this study were enriched with Paneth cells in both WT and *Atgl6l1*^ΔIEC^ organoids.

We then sought confirmation that *Atg16l1*^ΔIEC^ organoids were affected in their autophagy process. Consistent with the intestinal epithelial cell-specific *Atg16l1* deficiency, Western blot analysis confirmed *Atg16l1*^ΔIEC^ KO organoids were deficient in the Atg16l1 protein. Atg16l1 was detected in WT organoid samples at 68kDa, but not in *Atg16l1*^ΔIEC^ organoids even though a non-specific band is seen with the used antibody at 66kDa. In addition, we also observed LC3I to LC3II conversion in WT but not in the *Atg16l1*^ΔIEC^ organoids thus indicating that *Atg16l1* deletion leads to autophagy deficiency. As observed in previous studies, lack of *Atg16l1* resulted in impairment of autophagy as corroborated by reduced levels of LC3II (**Figure 2C**; Cadwell et al., 2008; Patel et al., 2013). Altogether these observations validate *Atg16l1*^ΔIEC^ organoids as a robust model for investigating the impairment of autophagy in epithelial homeostasis.

### Alteration in the proteomic abundance profiles upon autophagy-impairment

In order to determine the functional significance of the *Atg16l1* deficiency in Paneth cells, we established an integrated workflow (**Figure 1**) combining computational approaches to integrate and interpret the experimental readouts. We measured the protein levels in Paneth cell-enriched organoids derived from WT mice and mice harboring the *Atg16l1* deficiency with three biological replicates generated per condition tested. Our proteomic experiments detected 283 mouse proteins corresponding to 284 human ortholog proteins with altered abundances at the cut-offs used (p < 0.05, absolute relative fold change ≥ 2, number of unique peptides ≥ 2) (**Supplementary table 5**). Our initial functional analysis showed that proteins with altered abundance were related to at least 18 functional processes (**Figure 3**), and that the majority (70%) of all of these proteins were detected at levels twice greater than those found in WT organoids (**Supplementary table 6**), suggesting that the observed higher abundance could be a due to autophagy impairment.

**Figure 3.**
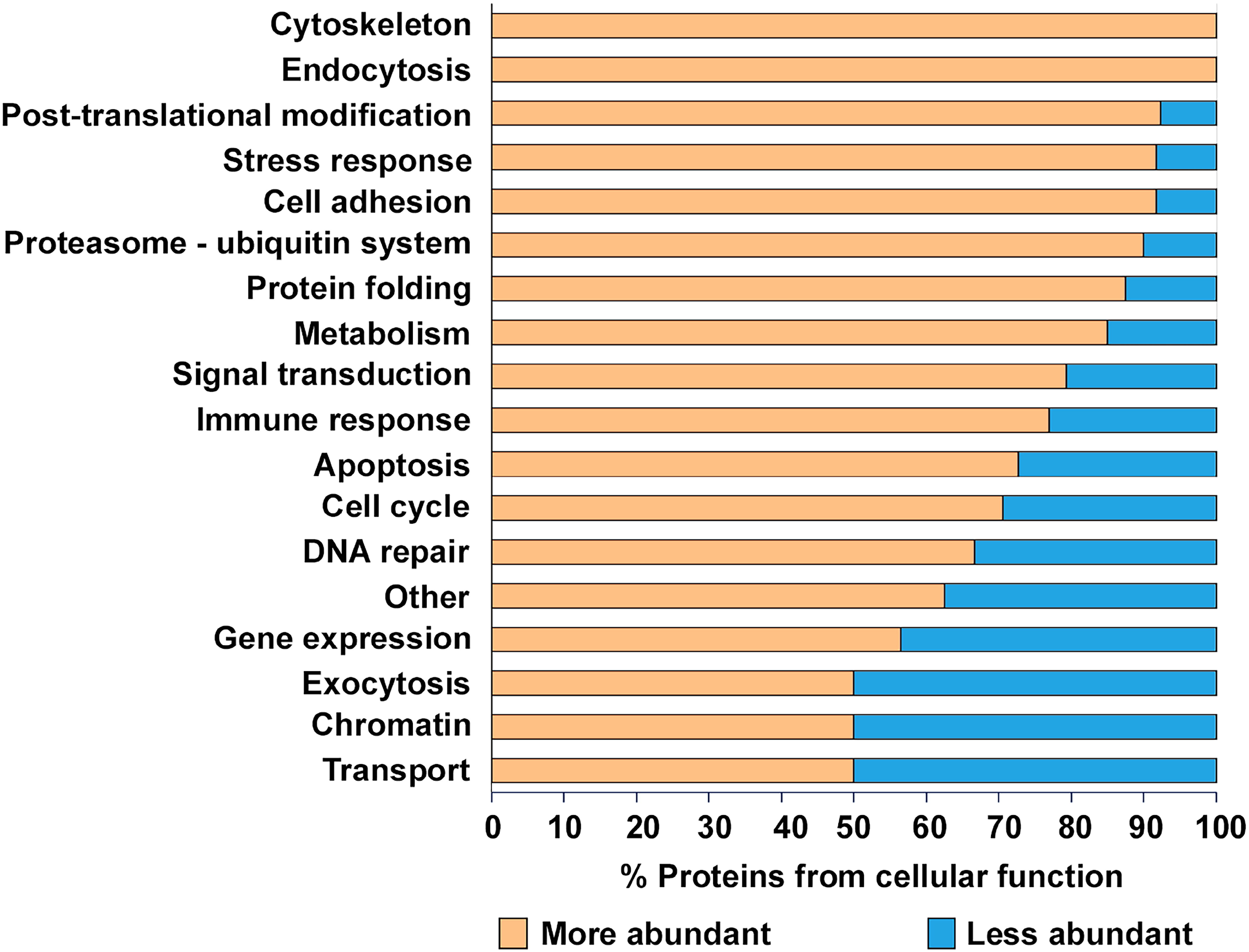
Percentage of higher and lower abundance proteins in different cellular functions. Proteins with higher abundance are marked with yellow and lower abundance proteins with blue background. “Other” category contains all of the proteins which have not fit to other functional groups.

### Proteins potentially targeted by selective autophagy have altered abundances in *Atg16l1*^ΔIEC^ Paneth cell organoids

Since the primary role of autophagy is to identify, target and recycle damaged proteins, altered protein levels in *Atg16l1*^⟀IEC^ Paneth cell-enriched organoids reflect the possible effect of a disrupted autophagy process. To determine whether autophagy directly or indirectly affects the proteins that are differentially abundant in the autophagy-impaired background, we compared the altered proteins with the target proteins of known selective autophagy receptors and adaptors, such as p62, LC3 and Atg16L1 (**Table 1**). This network analysis and the subsequent functional investigations were performed using human data (by inferring the human orthologs of the differentially abundant mice proteins) due to increased data availability on human networks/ontologies and thereby an increased coverage. By incorporating information about the binding partners (using experimental evidence and structure-based predictions) of the human orthologs, we identified the autophagy targeting proteins which could potentially target the altered proteins in Paneth cell organoids. 116 proteins (41%; P-value 0.049) with altered abundance in autophagy-impaired organoids and more importantly, 85 proteins with increased abundance (43.14%; P-value 0.043) (**Supplementary table 7**) were found to be potentially targeted by at least one of the three autophagy-related proteins (p62, LC3 and ATG16L1). This indicates that the proteins with increased levels in autophagy-impaired Paneth cell organoids are targeted for degradation by selective autophagy in normal organoids where autophagy is functional and not compromised. Overlap analysis of the altered proteins individually targeted by p62 (upregulated in *Atg16l1*-deficient organoids as expected (Ichimura et al., 2008), LC3 or ATG16L1 indicates that only a small proportion (19%) of the altered proteins potentially targeted by autophagy are targeted by more than one of the three autophagy proteins (**Figure 4, Supplementary table 8**). This suggests that the autophagy machinery potentially mounts a coordinated effort to specifically target certain groups of proteins.

**Figure 4.**
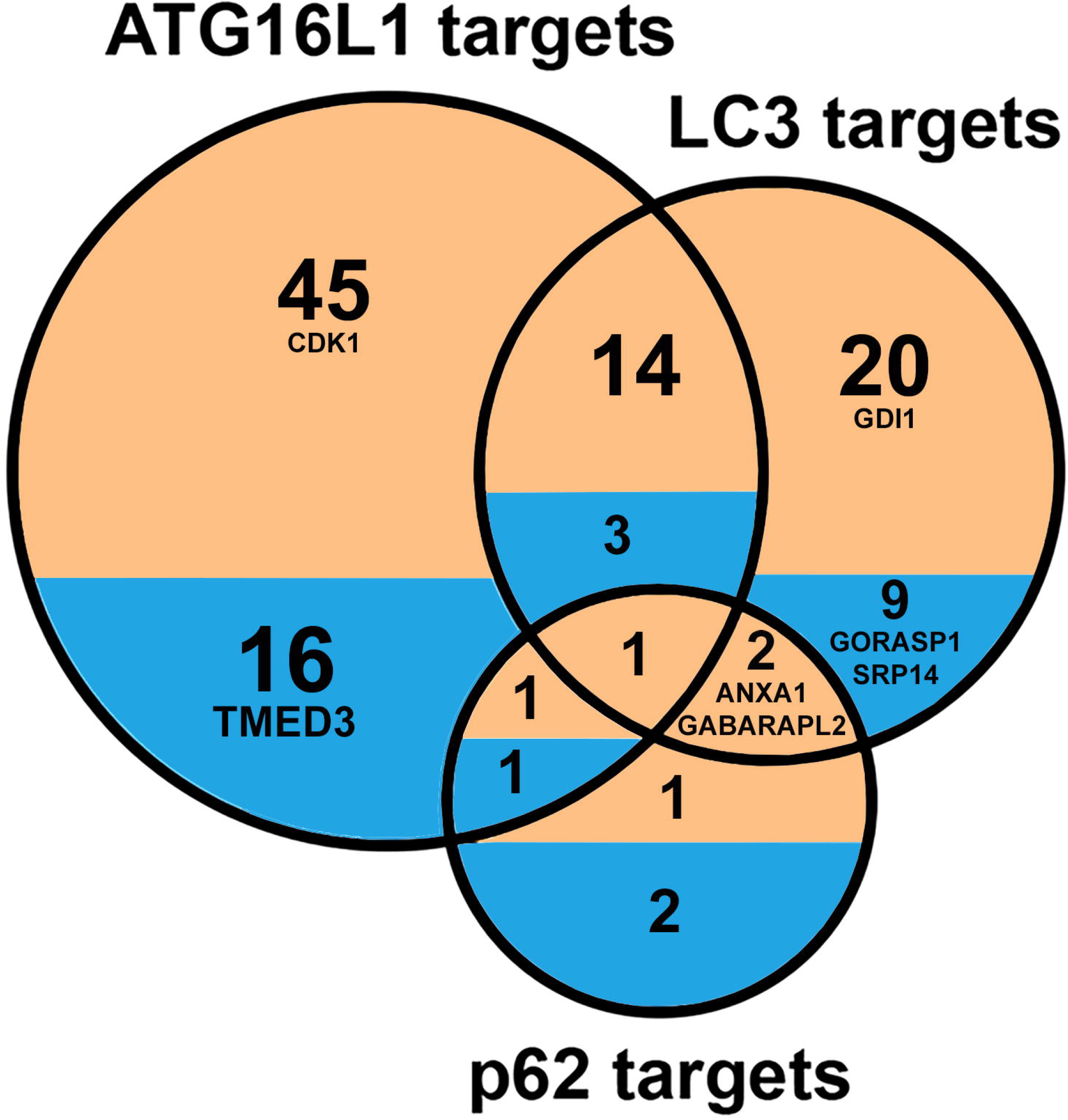
Overlap between the proteins with altered abundances in *Atg16l1*^⟀IEC^ Paneth cell organoids. Restricted to proteins potentially targeted by each of the three selective autophagy mediating proteins p62, LC3 and Atg16l1 under normal circumstances without any autophagy defects.

**Table 1.**
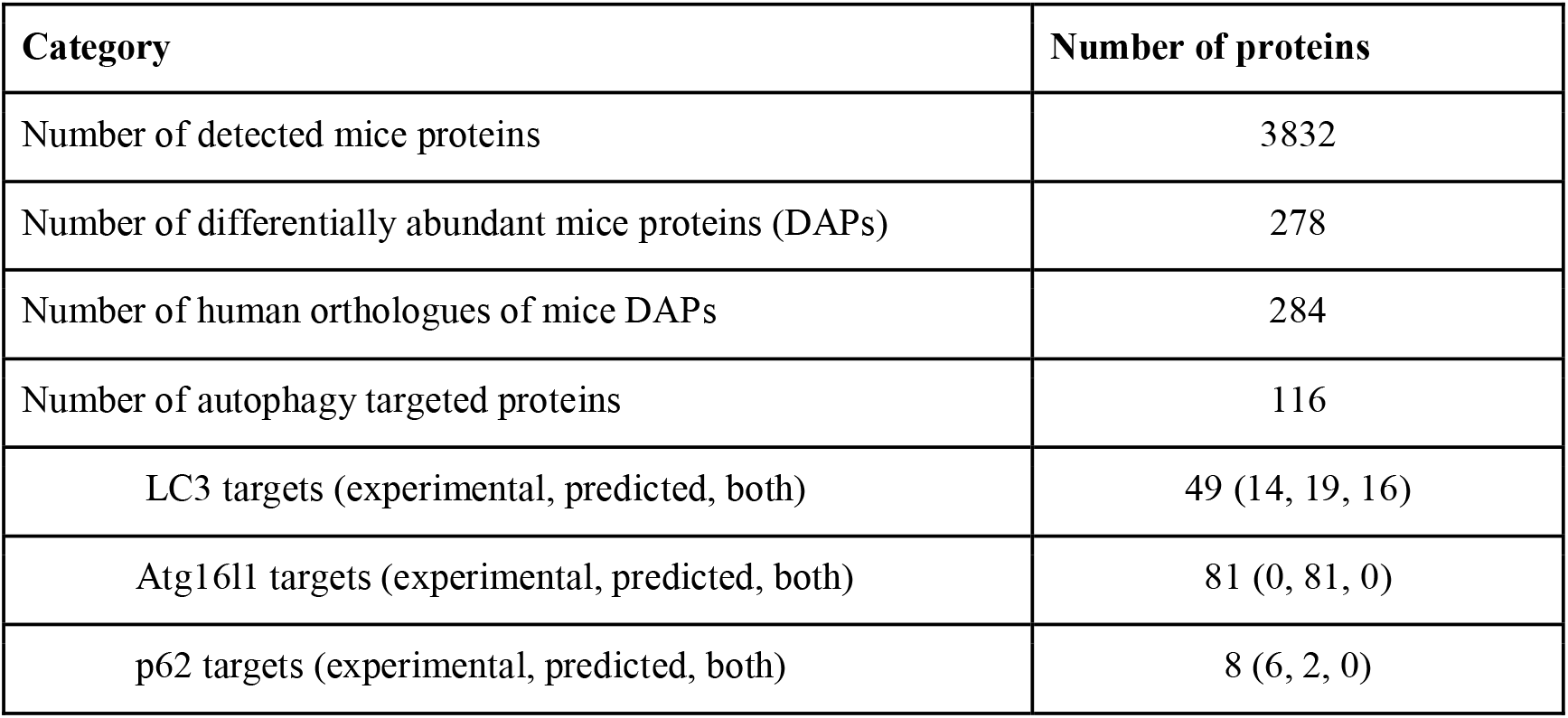
Effect of the *Atg16l1*^⟀IEC^ mutation on the alteration of protein abundances in Paneth cell-enriched organoids.

### Identification of Paneth cell functional processes affected by autophagy-impairment

To determine which cellular functions could be affected due to the altered protein abundances upon autophagy-impairment, we analysed the protein functions using manual curation of experimental evidence and Gene Ontology Biological Process terms (**Supplementary tables 9, 7**). We identified altered functional processes, such as apoptosis, exocytosis, DNA repair etc that could be dependent on autophagy-mediated protein degradation (**Figures 5, 6, S1 and S2; Supplementary tables 10**).

**Figure 5.**
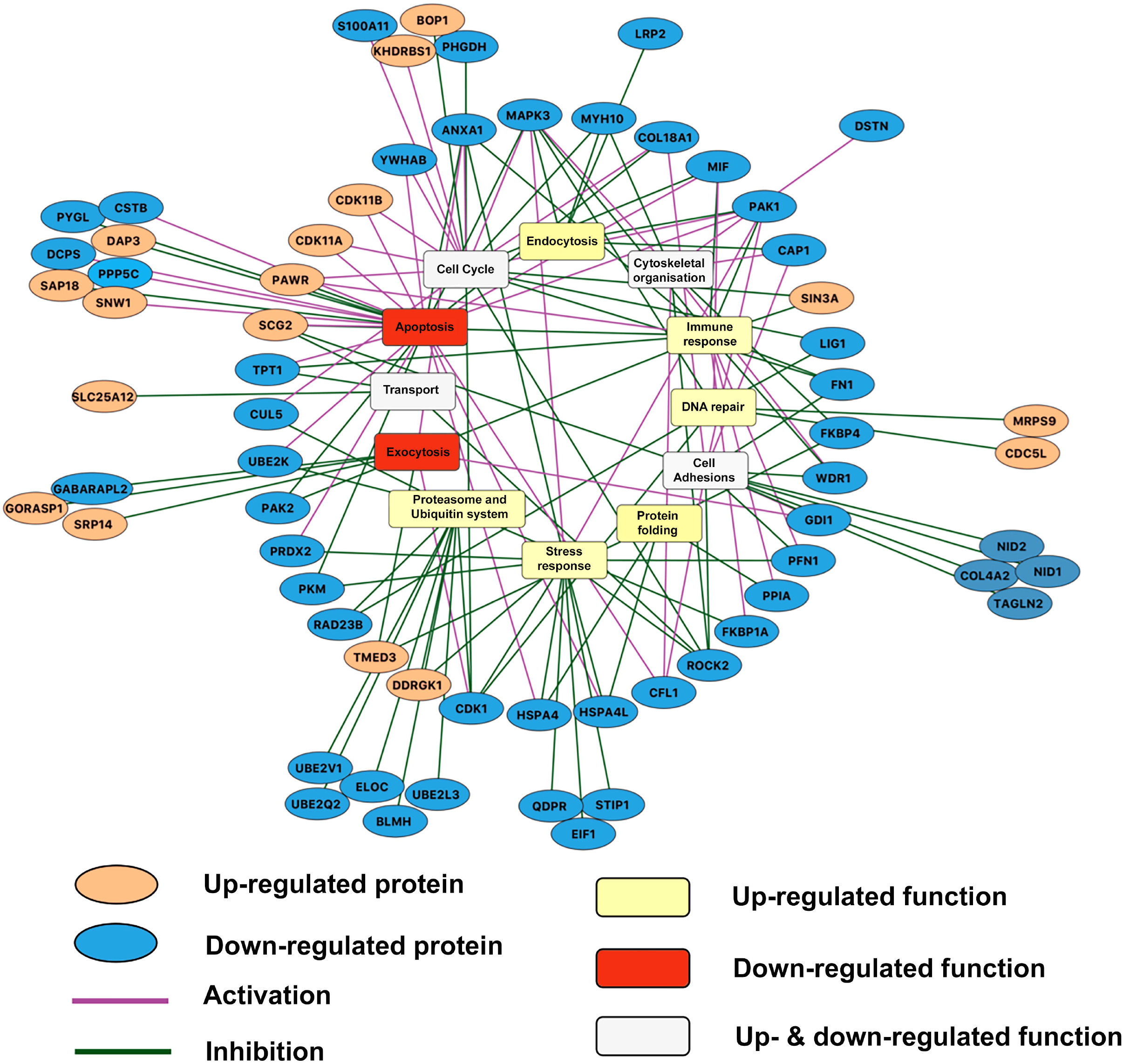
Potential autophagy dependency of the altered functional processes inferred from the proteomic profile of the *Atg16l11*^⟀IEC^ Paneth cell organoids using our integrated approach. The autophagy dependency of the proteins with altered abundances (orange ellipsoids for proteins with increased abundance, blue ellipsoids for proteins with decreased abundance) are represented highlighting the effect of proteins in the processes (purple edge for activation and green edge for inhibition) as well. The aggregated trends of the altered functional processes as determined by integrative approach (see Methods section) are indicated (yellow rectangles for up-regulated functional processes; red rectangles for functional processes; white rectangles for functional processes, which are both up- and down-regulated). Proteins outside from the circle are grouped to the process which they are involved in to increase the clarity of the figure. The figure was created using Cytoscape.

**Figure 6.**
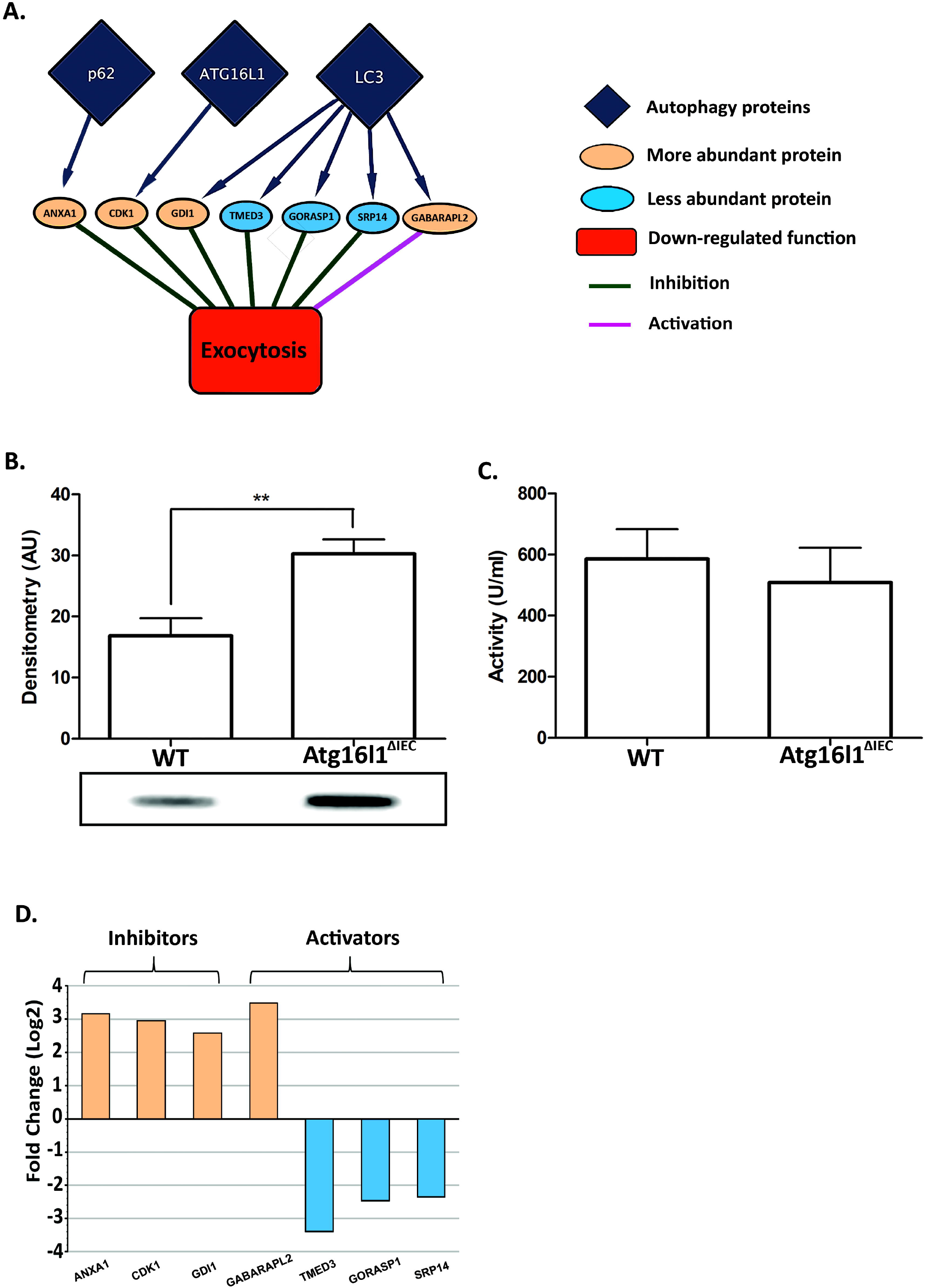
*Atg16l11*^⟀IEC^ deficiency in Paneth cell organoids and its impact on exocytotic proteins (n=3). (A) Proteins belonging to the functional category of exocytosis with lower abundance reflecting the impact of autophagy impairment on Paneth cell functions such as granule processing and release through exocytosis. (B) Western blot analysis for Paneth cell-derived lysozyme on cellular extracts from WT or *Atg16l11*^⟀IEC^ organoids expressing similar level of CD24 (**Figure 2B**). (C) Lysozyme activity measured in culture medium of 2D WT and *Atg16l11*^⟀IEC^ organoids as reporter of Paneth cell exocytosis. (D) Activators or inhibitors of exocytosis-related proteins found to be differentially abundant upon autophagy impairment. Blue and orange bars correspond to proteins with decreased and increased abundances, respectively. Overall these changes support the observed reduction of exocytosis.

Since post-translational regulators can elicit positive and negative effects on functional processes, we integrated an extensive literature curation evaluating the effect of each differentially abundant protein on associated functional processes (**Supplementary table 11**). For each functional process, we separately calculated an aggregated trend (see Methods for detail) to determine how the altered protein levels and the identified effect result in up- or down-regulation of process (**Table 2**). **Figure 5** outlines the potential autophagy-dependent functional categories that were altered and their aggregated trends. Overall, 66 differentially abundant proteins were described by functions. Interestingly, based on the aggregated trends, we observed that 14 of the 16 altered functional processes were either uniquely up-regulated or bi-directionally modulated (both up- and down-regulated), while only two functional processes were uniquely down-regulated (**Table 2**). This suggests that the overall consequence of autophagy-impairment in Paneth cells is predominantly characterized by the (over)activation of various functions. These include processes such as DNA repair, endocytosis, immune response and mitochondrial organization. Some of the up-regulated functions such as endocytosis and immune functions have previously been directly associated with autophagy (Deretic et al., 2013; Levine et al., 2011; Tooze et al., 2014). The two uniquely down-regulated functional processes, apoptosis and exocytosis, have also been associated with autophagy (Brooks et al., 2012; El-Khattouti et al., 2013; Tooze et al., 2014). We observed for example that 19/25 (76%) of apoptosis-related proteins that are more abundant when autophagy is impaired have an inhibitory impact on apoptosis, probably resulting in overall down-regulated apoptosis (**Supplementary Figure S1**). Similarly, 5/7 (>71%) of DNA repair-related proteins that are more abundant upon autophagy alteration have a activating impact on DNA repair (**Supplementary Figure S2**). Thus, taken as an extension to previous findings, our results show that autophagy-mediated protein degradation can regulate key Paneth cell functions, such as exocytosis, and potentially affect the activity of apoptosis regulators.

**Table 2.**
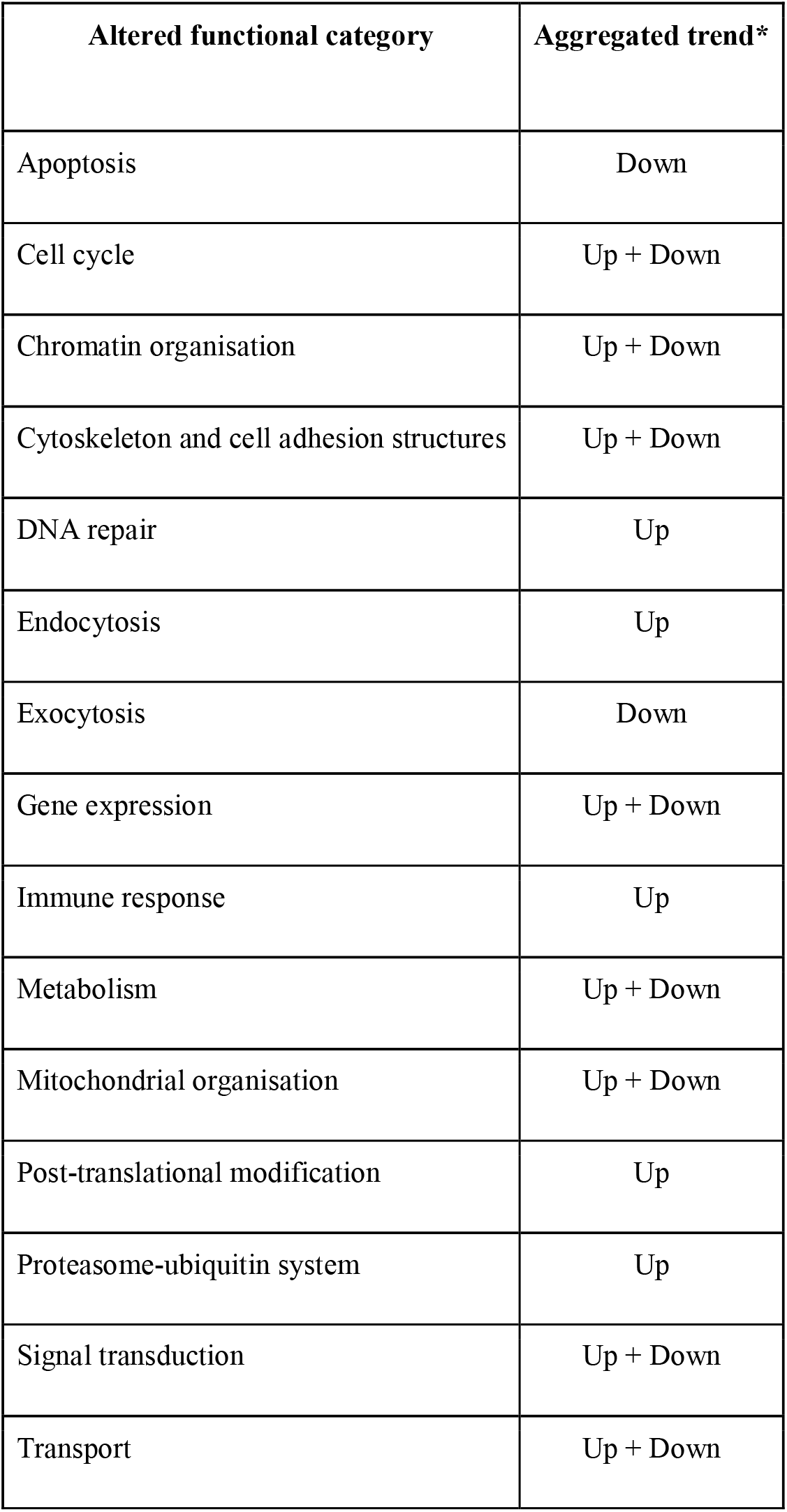
The aggregated trends of the functional categories which correspond both to the proteins with altered abundances and are potentially targeted by selective autophagy in response to the *Atg16l1*^⟀IEC^ mutation in Paneth cell-enriched organoids. * Aggregated trends were determined as described in the methods section.

### Validation of autophagy effect on protein degradation using transcriptomic data analysis

Change in abundance of proteins can be a consequence of altered gene expression, protein production and protein turnover. To validate that the observed difference due to autophagy impairment is mediated through protein turnover and not gene expression changes, we measured the transcriptomic profiles of Paneth cell-enriched organoids derived from mice lacking or not Atg16l1 in their intestinal epithelium. Comparing the FC value of genes and proteins which occurred in both proteomic and transcriptomic profiles can help to understand whether change in protein abundance depends on gene expression or on protein turnover. The genes coding for the 66 differentially abundant proteins that are predicted autophagy targets (LC3, p62 or ATG16L1 binding partners) were therefore analysed from transcriptomics data generated on additional organoid samples (**Supplementary table 12**). As a result, we found that 44 proteins, from the 66 differentially abundant candidates, were also differentially transcribed (**Supplementary table 12**). However, most of them (40 out of the 44 proteins) showed noticeable differences at both transcriptional and protein levels using the cutoff value defined in the Methods section: in every case, the fold change value for the transcriptomics data was much lower than the change in abundance of the protein expressed from the gene. Therefore, we assume that changes in protein abundance level happen due to the impaired autophagy-mediated protein degradation.

### Impact of autophagy-impairment on exocytosis in Paneth cells

As secretory cells, Paneth cells are reliant on high levels of protein biosynthesis and secretion, the latter being strongly reliant on functional autophagy. We assessed whether the autophagy impairment had an effect on the levels of proteins associated with exocytosis. Interestingly, our integrative analysis (using the workflow explained before) of the proteomic response revealed that exocytosis could be repressed in the absence of functional autophagy. **Figure 6A** shows the altered levels of exocytosis proteins in the Atg16l1^⟀IEC^ Paneth cell organoids as well as the autophagy targeting proteins, which could be modulating them. This result is in agreement with the already established importance of autophagy in the exocytosis-mediated secretion of antimicrobial peptides (AMPs) (Brooks et al., 2012; Cadwell et al., 2008; Cadwell et al., 2009a; Cadwell et al., 2009b; Gassler, 2017; Tschurtschenthaler and Adolph, 2018). We also determined experimentally that lysozyme levels detected within organoids were significantly greater when autophagy was impaired (**Figure 6B**) than in WT organoids. However, levels of lysozyme secreted into the culture medium were slightly reduced although not significantly different whether *atg16l1* was present or absent (**Figure 6C**), suggesting a defective exocytosis pathway upon impaired autophagy. Detailed analysis of our proteomic data showed that proteins targeted by LC3, Atg16l1 and p62 (**Figure 4**) and involved in the inhibition of exocytosis were found to be more abundant when autophagy was impaired. The opposite effect was observed with autophagy targeted proteins involved in the activation of exocytosis being less abundant upon autophagy-impairment (**Figure 6D**). This agrees with the negative alteration of exocytosis of AMPs that we observed in our validation assays (**Figure 6B, C**).

The exocytotic pathways facilitating the secretion of proteins are mediated either by general mechanisms involving the endoplasmic reticulum and the Golgi apparatus (Barlowe and Miller, 2013) in an autophagy-independent manner, or, in the case of antimicrobial proteins through the recently discovered LC3- and autophagy-dependent secretory process (Bel et al., 2017; Ponpuak et al., 2015). So far it was neither clear whether these two exocytotic pathways are co- or independently-regulated, nor whether they share some of the proteins involved and target overlapping proteins. Here, we revealed that proteins with exocytosis functions having higher abundance levels upon impaired autophagy could be potential autophagy-targets; these include SRP14 (Signal recognition particle 14 kDa protein), GORASP1 (Golgi reassembly-stacking protein 1) and TMED3 (Transmembrane emp24 domain-containing protein 3; **Figure 6D; Supplementary table 13**). This unexpected result raises the question whether to revisit the autophagy relatedness of the ER/Golgi-specific pathway, which shunts lysozyme into secretory granules involved in exocytosis. Based on these observations, it is plausible that autophagy could have a direct effect on processes, which are thought to be autophagy-independent.

## Discussion

Using a multidisciplinary combinatorial approach generating integrating interaction networks from proteomic data from murine Paneth cell-enriched organoids, interaction networks and validatory experiments, we revealed Paneth cell functional processes that are dependent on autophagy. Atg16l1 has been described as a pivotal autophagy protein in the last decade and it was shown that dysfunctional Atg16l1 leads to impaired formation of autophagosomes and poor degradation of long-lived proteins (Kaser and Blumberg, 2014; Saitoh et al., 2008). Our study focused on the role of autophagy in Paneth cell homeostasis, in particular on the consequences of impaired autophagy on Paneth cell functions. We used Paneth cell-enriched organoids derived from mice lacking the *Atg16l1* gene specifically in intestinal epithelial cells. This model may present the inconvenience of not being entirely composed of Paneth cells, and sorting Paneth cells from 3D organoids would have been an equally valid option, yet technically more challenging in view of generating sufficient amount of materials for proteomic analysis. Furthermore, it was recently shown that *in vitro* cultured organoids enriched for a specific cell type such as Paneth cells exhibit features that recapitulate better functions of *in vivo* Paneth cells than normally differentiated organoids (Mead et al., 2018). Using a *lyz-Cre* mouse model in future studies in combination with single cell transcriptome profiling will also confirm the impact of autophagy impairment we measured on Paneth cells.

Atg16l1 is an important component of the autophagy machinery whose human ortholog was previously associated to digestive pathologies such as Crohn’s disease. We determined quantitative proteomics profiles of Paneth cells-enriched organoids with functional or impaired autophagy. We developed and applied a computational systems biology approach based on the analysis of proteomics data generated from Paneth cells with functional and impaired autophagy and integrated multiple types of already existing but so far unconnected disparate information (protein-protein interaction networks, information about proteins known to be targeted by autophagy and functional information about proteins displaying differential abundance when autophagy was impaired). Integration of these data with the interaction networks of selective autophagy receptors and adaptors, such as p62, LC3 and ATG16L1, helped relating the degradation of the altered proteins to their regulation by autophagy. Furthermore, by incorporating known functions and biological processes attributed to the affected proteins, we identified various cellular processes, which could be dependent on a functional autophagy process.

As recently reported for stem cell-enriched organoids, our study emphasizes the robustness of systems-level approaches to fully capture the impact of major impaired cellular processes; in our case autophagy on homeostatic cellular functions (Lindeboom et al., 2018). The computational pipeline presented in this study enabled building regulatory networks of proteins displaying differential abundance upon autophagy impairment. To overcome the lack of mouse protein-protein interaction information involving the autophagy receptor and adaptor proteins, as well as to exploit the corresponding information already available in human datasets, we used the human orthologs of the mice proteins with altered abundances in mouse-derived *Atg16l1*^⟀IEC^ organoids. Although the cross-species extrapolation could be a source of uncertainties and possible missing information, the identified processes and their direction of modulation concur to a certain extent with already existing knowledge about the effects of autophagy-impairment. Other notable limitations of our study include the inability of the proteomic measurements to distinguish between the two isoforms of LC3 - LC3I and LC3II, thereby hindering interpretations about the role of the isoforms.

Strikingly, our analysis revealed that when autophagy is impaired upon lack of *Atg16l1*, nearly three hundred proteins display increased or decreased abundance, encompassing at least 18 functional processes (**Figure 3**). Transcriptomic analysis was carried out on Paneth cell-enriched organoids to identify the level of modulation of affected cellular processes. Most of the potential autophagy targeted proteins exhibited massive abundance discrepancies upon autophagy impairment but relatively small differential expression at the transcriptional level, confirming the strikingly stronger effect autophagy has on protein level regulation rather than on transcriptional regulation. Among the altered proteins, several had previously been associated with pathologies affecting Paneth cells, such as ANXA1 and FGA, which were previously reported to be altered in inflamed mucosal tissue or epithelial cells from Crohn’s disease patients (Barceló-Batllori et al., 2002; Iskandar and Ciorba, 2012; Meuwis et al., 2007). We observed that when autophagy is impaired in Paneth cells, most of the differentially abundant proteins are present in greater abundance than in normal Paneth cells, thus suggesting that degradation through autophagy plays a key role in maintaining the intracellular concentrations of these proteins.

The developed computational pipeline enhances our understanding about the underlying mechanisms involved in autophagy-mediated degradation by integrating multiple levels of information such as protein-protein interactions, autophagy mediated selective protein degradation, inhibitory/stimulatory relationships between the altered proteins and functional processes. Notably, to reduce the impact of linear assumptions in interpreting the impacts of proteomic changes on functional processes, we determined the aggregated trends for the functional processes by incorporating the direction of protein level alterations and the stimulatory/inhibitory relationships between the altered proteins and the functions. Furthermore, by bringing together different levels of information, our approach helps explain the mechanistic underpinnings between the processes corresponding to the proteins with altered abundances and autophagy. Capturing such process level dependencies on cellular autophagy and their modulation would be difficult by using singular levels of information in isolation. For example, Zhang *et al* measured the proteome level changes in primary human fibroblasts which were impaired in autophagy as a means to explain the purported dependency of protein degradation on macroautophagy (Zhang et al., 2016). Patella *et al* identified proteomic alterations under conditions of autophagy blockage in endothelial cells to explain the potential role of autophagy in maintaining endothelial permeability (Patella et al., 2016). Similarly, various other studies have profiled the global proteomic changes in response to artificial impairment of autophagy by knocking out critical autophagy genes (Avin-Wittenberg et al., 2015; Mathew et al., 2014). Studies combining different -omic readouts have also been performed in various contexts to understand the role of autophagy in various processes and phenotypes (Chen et al., 2017; Kramer et al., 2017; Masclaux-Daubresse et al., 2014; Stingele et al., 2012). However, the studies aforementioned do not provide an explanation as to how the proteome level alterations are indeed dependent on autophagy from a mechanistic point of view. In this study, using networks and integration of heterogeneous datasets, we provide information on new mechanisms by which several cellular processes such as exocytosis, DNA repair and apoptosis are dependent on autophagy.

Upon microbial invasion or inflammation-mediated cellular damage, cells respond by activating apoptotic cell death. In general, autophagy and apoptosis are negatively correlated under most homeostatic conditions (Mariño et al., 2014), although altered cellular settings can drive autophagy to lead to programmed cell death. The interactions between autophagy and apoptosis processes are highly complex (Gump and Thorburn, 2011; Mariño et al., 2014). Interestingly, our observations showed a positive correlation between autophagy and apoptosis (apoptosis being inhibited in the autophagy-impaired organoids; **Supplementary Figure 1**). When autophagy is impaired, the observed down-regulation of apoptosis could prevent the perturbed Paneth cells from sacrificing themselves, which would then be compensated for by outcomes such as upregulation of DNA damage repair functions as suggested previously (Basu and Krishnamurthy, 2010; Nowsheen and Yang, 2012; **Supplementary Figure 2**). However, further experiments are needed to confirm the assumption about the role of DNA repair and apoptosis in Paneth cells and how the disruption of these processes could contribute to the pathogenesis of impaired autophagy-associated diseases.

ATG16L1 is not only known to be required for the normal functioning of autophagy but also has physiological relevance. The intestinal epithelium in patients with inflammatory digestive disorders is characterized by a prolonged period of stress as a result of chronic inflammation and malfunction of antimicrobial innate defences. This is reflected in particular by the alteration of exocytosis of antimicrobials as well as the manipulation of the genetic/epigenetic machinery and organelles by invading pathogens and various other causative agents (Fofanova et al., 2016; Sartor, 2006). In this study, the downregulation of exocytosis that we observed in autophagy-impaired organoids was illustrated by the lysozyme accumulation within organoid cells and the consequential alteration of lysozyme secretion into the extracellular milieu. These results support our computational analysis and concur with the previously reported autophagy dependency of exocytosis (Brooks et al., 2012; Cadwell et al., 2008; Cadwell et al., 2009a; Cadwell et al., 2009b; Gassler, 2017; Tschurtschenthaler and Adolph, 2018) (**Figures 4, 6**). Impaired autophagy can therefore have dramatic consequences on innate defense mechanisms against microbial invasion of the gut epithelium by deregulating the protein degradation of key exocytotic proteins.

The Paneth cell-enriched organoids we derived from the *Atg16l1*^⟀IEC^ mouse model could be perceived as a biased representation of the role autophagy impairment has in inflammatory diseases (infectious or chronic), as it does not consider other intrinsic factors such as mutations in non-autophagy related genes which have been shown to contribute to those pathologies (e.g. proteins like PARP-2, IFI35, S100A12, CRP and S100A8 previously shown to contribute to IBD pathogenesis (Cheluvappa et al., 2014; de Jong et al., 2006; Jijon et al., 2000; Leach and Day, 2006; Vermeire et al., 2006). These biomarkers are indicative of an inflammatory state, some of them are mostly detected in the serum, but indicators detected in faecal samples predict more accurately the state of digestive inflammatory state. For example, inhibition of PARP dampens inflammation associated with colitis (Jijon et al., 2000), while elevated levels of CRP, S100A8 for example have been associated with inflammatory pathologies. Yet in our study, these proteins were not found differentially abundant when autophagy was impaired, or were fluctuating in the opposite direction. These discrepancies could reflect the differences in the sample type as these studies did not focus on Paneth cells only. Complementary experiments and predictions would nonetheless help highlight some of the aspects of the molecular regulatory mechanisms that contribute to the pathogenesis of digestive diseases upon alteration of autophagy.

Our integrative analysis not only captured already known phenomena, namely the autophagy dependency of exocytotic functions associated with granule release, but also highlighted that the degradation regulation of many differentially abundant proteins, including those of exocytosis proteins occurring in an autophagy-dependent manner. The presented study therefore extended the list of proteins for which the degradation rate was already known to be regulated by autophagy. More interestingly, our analysis revealed additional cellular processes which could mediate the effects of autophagy impairment on Paneth cell functions. Taken together and using a mouse model where autophagy is impaired, and organoids to capture Paneth cells, we identified various cellular processes which are dependent on autophagy and the failure of which further could contribute to the pathogenesis of major digestive pathologies.

## Acknowledgements

The authors are grateful for the helpful discussions to the past and present members and visitors of the Korcsmaros, Hall and Wileman groups.

## Author contributions

EJ and ZM performed the experimental work with the organoids. LG, PS, AT and TWr carried out the bioinformatic analysis. DD, JB, MJ, UM created and tested the mice model, SA carried out the proteomics study. IH performed the lysozyme assays. AW, PP, SC, LJH, WH, FDP, TWi, TK designed and supervised the experiments. EJ, LG, PS, AT, IH and TK wrote the manuscript.

---

**Supplementary figure 1. Alterations describing the modulation of apoptosis in response to the *Atg16L1*^⟀IEC^ mutation**.

**Supplementary figure 2. Alterations describing the modulation of DNA repair in response to the *Atg16L1*^⟀IEC^ mutation**.

**Supplementary table 1. Specific primers used in the study**.

**Supplementary table 2. Discarded biological process terms not related to intestinal functions**.

**Supplementary table 3. Effect of the *Atg16L1*^⟀IEC^ mutation on protein abundances in normal organoid control experiment**.

**Supplementary table 4. Effect of the *Atg16L1*^⟀IEC^ mutation on protein abundances in Paneth cell-enriched organoid control experiment**.

**Supplementary table 5. Post-processed results from the measurements comparing the proteomic profiles of *Atg16L1*^⟀IEC^and WT Paneth cell-enriched organoids**.

**Supplementary table 6. Human orthologs of the mouse proteins with altered abundances**.

**Supplementary table 7. Evidence suggesting the targeting of proteins with altered abundances by the autophagy proteins (p62, LC3, ATG16L1**).

**Supplementary table 8. Summary of abundance changes of proteins potentially targeted by autophagy and their assigned gene ontology terms**.

**Supplementary table 9. Functional evidence indicating the stimulatory/inhibitory effect of the proteins with altered abundances on their corresponding processes**.

**Supplementary table 10. The aggregated trends of the gene ontology terms corresponding to the altered proteins in Paneth cells upon the deletion of *Atg16l1***.

**Supplementary table 11. Functional processes and their aggregated trends in Paneth cells upon the deletion of *Atg16l1***.

**Supplementary table 12. Overlapping functional proteins and their FC value in proteomics and transcriptomics studies**

**Supplementary table 13. Exocytotic proteins found to be differentially abundant upon autophagy impairment**.

