## Supplementary table 1 for "Integrative analysis of Paneth cell proteomic and transcriptomic data from intestinal organoids reveals functional processes dependent on autophagy"

Genotype/Gene	Forward Primer Sequences	Number of qPCR cycles

Reverse Primer Sequences

*atg16l1 ^Fl/+^*	CTGAACAGTTAAGTTCCTAG	N/A

CCAAGAGACACTGACATAGG

*atg16l1 ^Fl/- Vil-Cre^*	GACGGAAATCCATCGCTCGACCAG	N/A

GACATGTTCAGGGATCGCCAGGCG

*Villin*	GTGGAATGGGCCAGAGAGT	20

CATAATTGCCATCAGCTGTGG

*cd24*	TTGCTGCTTCTGGCACTGC	20

GGAGACCAGCTGTGGACTGC

*chromogranin A*	CCAGTTCCCACTTCCATGC	27

CCTTCAGACGGCAGAGCTTC

*lgr5*	ATACCGGAGCGAGCGTTC	25

TGGCAGTTCCTGTCAAGTGA

*muc2*	GTTGCTCAATGAGATGGAGGT	20

AAGCTGCATGACTGGAAGC

*β-actin*	GAGGCCCCCCTGAACCCTAAG	20

GAACCGCTCGTTGCCAATAG
