## Supplementary figures and images for "Integrative analysis of Paneth cell proteomic and transcriptomic data from intestinal organoids reveals functional processes dependent on autophagy"

### Supplementary figure 1

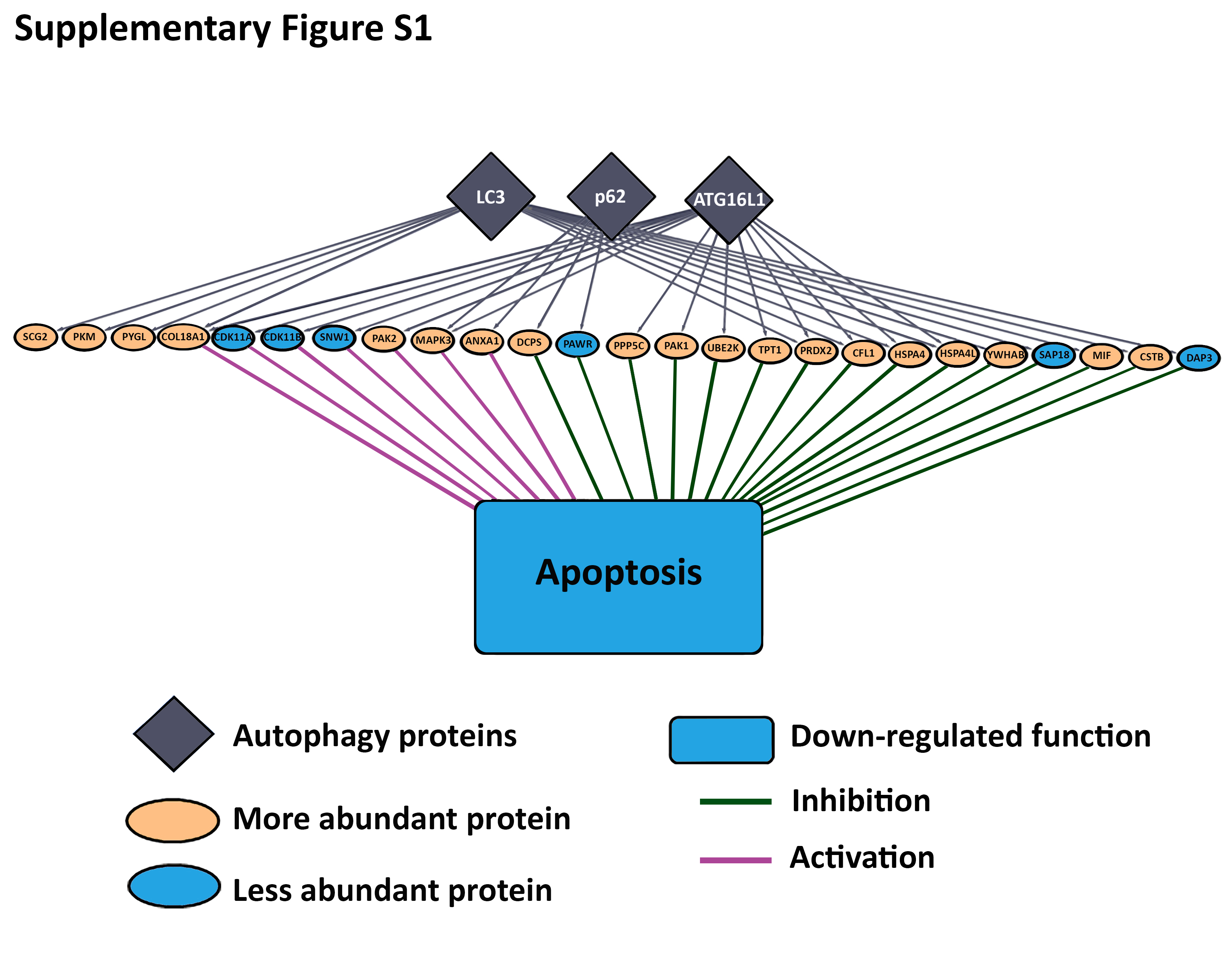

### Supplementary figure 2

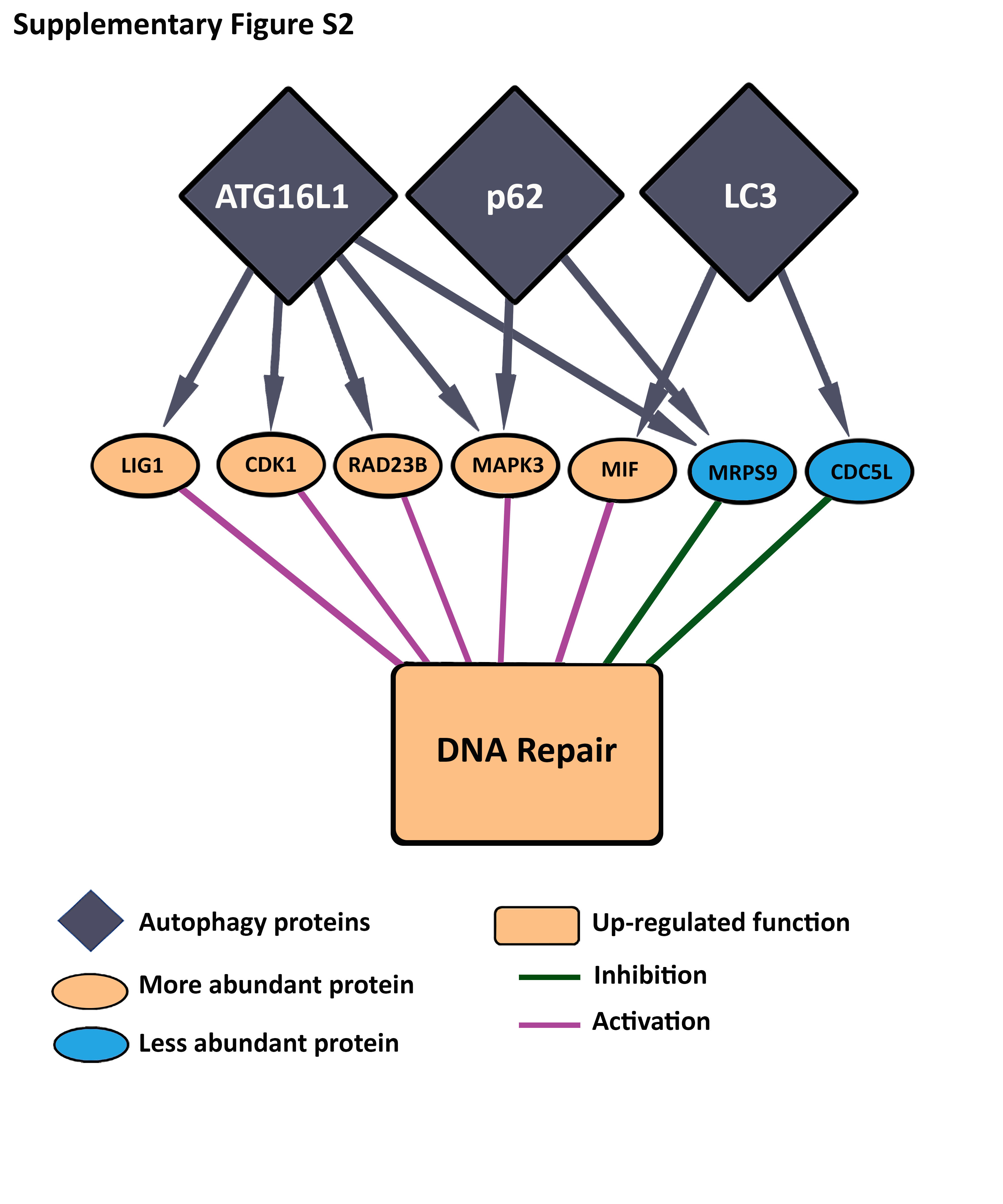
