## Supplementary table 13 for "Integrative analysis of Paneth cell proteomic and transcriptomic data from intestinal organoids reveals functional processes dependent on autophagy"

| **Gene symbol** | **Trend in alteration (absolute fold change)** | **Activator/Inhibitor** | **Function** | **Reference** |
| --- | --- | --- | --- | --- |
| **ANXA1** | **UP (3.17)** | **Inhibitor** | **Inhibitor of hormone exocytosis** | **McArthur et al, 2009** |
| **CDK1** | **UP (2.96)** | **Inhibitor** | **Inhibits the assembly of the GM130-p115-giantin tether and thus the fusion of COPI vesicles** | **Wang et al, 2008** |
| **GDI1** | **UP (2.58)** | **Inhibitor** | **Recycling of proteins from their target membranes back to their vesicular pools** | **Garrett et al, 1994** |
| **GABARAPL2** | **UP (3.5)** | **Activator** | **LC3 (containing GABARAPL-2 protein) contributes to the envelopment and exocytosis of viruses during lytic infection** | **Cadwell and Debnath, 2017** |
| **TMED3** | **DOWN (3.42)** | **Activator** | **Bind to both COPI and COPII proteins and likely function in anterograde and retrograde transport between the ER and the Golgi** | **Jerome-Majewska et al, 2010** |
| **GORASP1** | **DOWN (2.48)** | **Activator** | **COPII vesicle coating, ER to Golgi vesicle-mediated transport** | **Gene Ontology** |
| **SRP14** | **DOWN (2.37)** | **Activator** | **Targeting secretory proteins to the rough endoplasmic reticulum membrane** | **Gene Ontology** |
