## Supplementary material for "Integrative analysis of Paneth cell proteomic and transcriptomic data from intestinal organoids reveals functional processes dependent on autophagy": Table 1

| **Category** | **Number of proteins** |
| --- | --- |
| Number of detected mice proteins | 3832 |
| Number of differentially abundant mice proteins (DAPs) | 278 |
| Number of human orthologues of mice DAPs | 284 |
| Number of autophagy targeted proteins | 116 |
| LC3 targets (experimental, predicted, both) | 49 (14, 19, 16) |
| Atg16l1 targets (experimental, predicted, both) | 81 (0, 81, 0) |
| p62 targets (experimental, predicted, both) | 8 (6, 2, 0) |
