## Supplementary material for "Integrative analysis of Paneth cell proteomic and transcriptomic data from intestinal organoids reveals functional processes dependent on autophagy": Table 2

| **Altered functional category** | **Aggregated trend*** |
| --- | --- |
| Apoptosis | Down |
| Cell cycle | Up + Down |
| Chromatin organisation | Up + Down |
| Cytoskeleton and cell adhesion structures | Up + Down |
| DNA repair | Up |
| Endocytosis | Up |
| Exocytosis | Down |
| Gene expression | Up + Down |
| Immune response | Up |
| Metabolism | Up + Down |
| Mitochondrial organisation | Up + Down |
| Post-translational modification | Up |
| Proteasome-ubiquitin system | Up |
| Signal transduction | Up + Down |
| Transport | Up + Down |
